# Human placental bed transcriptomic profiling reveals inflammatory activation of endothelial cells in preeclampsia

**DOI:** 10.1101/2021.10.18.464811

**Authors:** Laura Brouwers, Judith Wienke, Michal Mokry, Peter GJ Nikkels, Tatjana E. Vogelvang, Arie Franx, Femke van Wijk, Bas B. van Rijn

## Abstract

**Rationale:** Functional characteristics of endothelial cells (ECs) within the human placental bed are unknown and may provide insight into the adaptive biology of ECs in disorders of vascular remodelling like preeclampsia.

**Objective:** To determine transcriptional profiles of human placental bed ECs and systemic biomarker profiles in women with normal pregnancy, and women with preeclampsia, a condition characterized by extensive EC dysfunction, poor development of spiral arteries underlying the placenta and long-term susceptibility to atherosclerosis and hypertension.

**Methods & results:** We obtained biopsy samples from the uterine placental bed, of five women with preeclampsia with fetal growth restriction (FGR) due to impaired spiral artery development and four controls undergoing Caesarean section. CD31^+^CD146^+^ ECs were isolated and sorted by flow cytometry for RNA-sequencing using CEL-Seq2 protocol. Data were analyzed by unsupervised clustering, gene set enrichment (GSEA) and pathway analysis. 67 circulating biomarkers of EC function and inflammation were measured in 20 women with preeclampsia with FGR and 20 controls by multiplex immunoassay. Transcriptional profiling showed various differentially expressed genes (FDR<0.05) in placental bed ECs of preeclampsia patients, with enhanced activity of pathways associated with vasoconstriction, platelet activation and innate immunity. GSEA was suggestive of a VEGF- and PlGF deprived state of preeclampsia-derived ECs. Moreover, the transcriptomic profile was similar to that of human umbilical vein endothelial cells (HUVECs) treated with plasma from preeclampsia patients, pointing towards a central role for circulating factors in EC dysfunction. Unsupervised clustering of subjects by EC-related circulating factors identified distinct profiles for healthy pregnancy and preeclampsia, in particular for those women with low platelets and elevated liver enzymes, which was predominantly driven by sFLT-1, endoglin, PlGF, leptin, SAA-1 and sICAM-1.

**Conclusions:** We revealed inflammatory activation of EC and a key role for systemic factors in EC dysfunction in women with preeclampsia associated with impaired spiral artery development.

## INTRODUCTION

Preeclampsia is a serious multisystem vascular disorder characterized by new-onset hypertension and proteinuria, as well as vascular impairment of many organ systems, e.g. the liver, kidneys, and brain. Preeclampsia affects about 1-5% of women in the second half of pregnancy. Endothelial cell dysfunction is generally acknowledged as its key pathophysiological feature.(1) Severe, or early onset, preeclampsia is often associated with fetal growth restriction (FGR) as a result of insufficient placental function due to underdevelopment of the maternal arteries supporting placental growth.(2, 3) Women who experience preeclampsia are at an increased risk of arterial disease, including atherosclerosis, ischemic heart disease and stroke later in life, and preeclampsia is now considered as one of the strongest sex-specific risk factors for major cardiovascular events in women of a young age.(4, 5) The cause of preeclampsia and the sequence of events leading to the disease is unclear. However, impaired physiological transformation of the uterine vascular bed underlying the placenta (i.e. the placental bed), known as spiral artery remodeling, is thought to be the essential first stage preceding the symptomatic phase of generalized endothelial dysfunction in most cases.(6) Spiral artery remodeling is necessary for adequate blood flow to the developing placenta and fetus, and is histologically characterized by reorganization of vascular wall components including ECs, smooth muscle cells and the elastic lamina and new formation of a fibrinous layer within the vessel wall. It has been suggested that intramural invasion and EC replacement in response to migration of extravillous trophoblasts into the placental bed is a key mechanism for successful remodeling of spiral arteries.(7) In preeclampsia and FGR, spiral artery remodeling is often impaired or absent, leading to altered blood flow to the placenta. In spiral arteries with defective remodeling, histological signs of loss of EC function have been observed, e.g. thrombotic lesions and infiltration of foam cells and lipid deposits into the intimal and medial layer termed ‘acute atherosis’.(8–11) It has been hypothesized that these processes at the maternal-fetal interface, the generalized maternal endothelial dysfunction, and subsequent hypertension of preeclampsia are linked and result from the release of inflammatory and antiangiogenic factors into the maternal circulation.(12) Similar to other arterial disorders, e.g. during the early stages of atherosclerosis, women with preeclampsia consistently show increased activation of the systemic inflammatory response both during pregnancy and in the non-pregnant state, including elevated levels of C-reactive protein (CRP), interleukin (IL)-6 and IL-8.(13) In addition, endothelial cell activation is suggested by elevated levels of several endothelial and vascular cell activation molecules, i.e. E-selection and sVCAM-1.(14) Specific to preeclampsia are high levels of the anti-angiogenic factors sFLT-1 and endoglin and low levels of the pro-angiogenic agent PlGF, which are released in abundance by the placenta, and are thought to contribute to impaired EC function by sustaining an anti-angiogenic environment.(5, 15) Additionally, the complex homeostatic balance of vascular tone, which is normally maintained by the endothelium, is disturbed, leading to dysregulated and elevated blood pressure.(16, 17) ECs exposed to serum from women with preeclampsia show signs of dysfunction in vitro, indicating the presence of circulating factors involved in systemic endothelial dysfunction.(18, 19) Furthermore, women with preeclampsia have an increased risk of future cardiovascular disease, in particular for atherosclerosis, implying that a vulnerable maternal vascular constitution may contribute to both disorders by an increased susceptibility to EC dysfunction independent of pregnancy.(4, 11, 20)

Due to the practical and technical challenge of getting access to tissue from the human placental bed and isolating tissue-specific ECs from human tissue for reliable transcriptomic analysis, little is known about the functional changes of ECs at the maternal-fetal interface during preeclampsia. Here, we applied flow cytometry-assisted cell sorting of ECs for RNA-sequencing to investigate preeclampsia-induced changes in the transcriptomic profile of ECs within the human placental bed. We applied cluster and pathway analyses to identify functional changes in ECs associated with abnormal spiral artery remodeling typical of preeclampsia and FGR, in addition to circulating markers representative of systemic inflammation and endothelial activation by multiplex immunoassay.

## METHODS

### Patient selection and definitions

This study was part of the Spiral Artery Remodeling (SPAR) study. SPAR is a prospective multicenter study investigating spiral artery remodeling and pathology in women with and without preeclampsia and/or fetal growth restriction (FGR). Detailed description of the study design and sampling protocol was previously published.(11) In short, all women with a clinical indication for a Caesarean section for either preeclampsia or FGR, or both, were asked to participate in the study. In addition, women who delivered by primary elective Caesarean section after an uneventful pregnancy and without any major underlying pathology were enrolled as controls. We defined preeclampsia according to the most recent definition of the International Society for the Study of Hypertension in Pregnancy, as new-onset hypertension (≥140/90mmHg) after 20 weeks of gestation in combination with significant proteinuria (≥300mg/24h or protein/creatinine ratio ≥0.3 mg/mg), maternal organ dysfunction (i.e. renal insufficiency, liver involvement, neurological or hematological complications) or the presence of FGR.(21) Preeclampsia complicated by HELLP syndrome, which is considered as a more severe form of the same condition, was defined according to the presence of two or more of the following criteria: hemolysis (defined as serum lactate dehydrogenase (LDH) >600 U/L and/or haptoglobin 0.3 g/L), elevated liver enzymes (serum aspartate aminotransferase (AST) >50 U/L and/or serum alanine aminotransferase (ALT) >50 U/L), and a low platelet count (<100×10^9^/L).(22) FGR was defined as an ultrasonographical estimated fetal weight or abdominal circumference below the tenth percentile or a reduction in the standardized growth curves of ≥20 percentiles.(23) Placental insufficiency had to be the suspected cause of preeclampsia and FGR, and cases with confirmed chromosomal and/or congenital abnormalities were excluded. Further details and definitions used in the study can be found in our previous publication.(11)

For this study we chose to include women with severe disease on the basis of early-onset, i.e. onset and delivery of preeclampsia before 34 completed weeks of gestation, and the presence of FGR as confirmation of the placental origin of the disease. For EC transcriptomics we included only primiparous women, N=5 with preeclampsia, and N=4 women with healthy pregnancies. For the multiplex immunoassay, we included 20 patients with severe preeclampsia and FGR with histologically confirmed spiral artery pathology, and 20 healthy women with uneventful pregnancies with histologically confirmed normal spiral artery remodeling as controls. All patients were delivered by elective Caesarean sections, without any signs of labour, e.g. contractions or rupture of membranes. All patients provided written informed consent prior to participation. This study was reviewed and approved by the Institutional Ethical Review Board of the University Medical Center Utrecht, protocol reference number: 16-198 and was prospectively monitored for any adverse events.

### Placental bed biopsies

After delivery of the neonate and the placenta, as per routine procedures, the placental bed was manually located and two biopsies of the central placental bed from the inner uterine myometrial wall were obtained according to a pre specified protocol published previously.(11) In addition to the placental bed site, biopsies were taken from the incision site when the placenta was not situated on this part of the uterine wall.

### Isolation of placental bed ECs

The biopsy samples were collected in medium consisting of RPMI 1640 (Gibco) supplemented with Penicillin/Streptomycin (Gibco), L-glutamine (Gibco) and 10% fetal calf serum (FCS, Biowest) and minced into pieces of 1 mm^3^ in PBS (Gibco). The biopsies were enzymatically digested with 1 mg/mL collagenase IV (Sigma) in medium for 60 minutes at 37°C in a tube shaker under constant agitation at 120 rpm. To dissolve the remaining biopsy pieces after digestion and remove any remaining lumps, the biopsies were pipetted up and down multiple times and poured over a 100 μm Cell Strainer (BD Falcon). Cells were subsequently washed in staining buffer consisting of cold PBS supplemented with 2% FCS and 0.1% sodium-azide (Severn Biotech Ltd.) and filtered through a 70 μm cell strainer. For FACS sorting, the cells were incubated with surface antibodies against CD45, CD31 (PECAM-1), CD146 (MCAM), CD54 (ICAM-1), CD144 (VE-cadherin), CD105 (Endoglin), and CD309 (VEGFR2) (Supplementary Table 1) for 20 minutes in staining buffer at 4°C, washed in the same buffer and filtered through a 50 μm cell strainer (Filcon, BD). 2000 cells of the CD45^−^CD31^+^CD146^+^ cell population were sorted into eppendorfs containing 125 μL PBS on one of the two available FACSAria™ II or III machines (BD). After sorting, 375 μL Trizol LS (Thermo Fisher Scientific) was added to each vial and vials were stored at −80°C until RNA isolation.

### RNA isolation

For RNA isolation, vials were thawed a room temperature and 100 μL chloroform was added to each vial. The vials were shaken well and spun down at 12000xg for 15 minutes at 4°C. The aqueous phase was transferred into a new tube and RNA was mixed with 1ul of GlycoBlue (Invitrogen) and precipitated with 250 μL isopropanolol. Cells were incubated at −20°C for one hour and subsequently spun down at 12000xg for 10 minutes. The supernatant was carefully discarded and the RNA pellet was washed twice with 375 μL 75% ethanol. Vials were stored at −80°C until library preparation.

### Whole transcriptome sequencing and data analysis

Low input RNA sequencing libraries from biological sorted cell population replicates were prepared using the Cel-Seq2 Sample Preparation Protocol(24) and sequenced as 2 x 75bp paired-end on a NextSeq 500 (Utrecht Sequencing Facility). The reads were demultiplexed and aligned to human cDNA reference using the BWA (0.7.13).(25) Multiple reads mapping to the same gene with the same unique molecular identifier (UMI, 6bp long) were counted as a single read. RNA sequencing data were normalized per million reads and differentially expressed genes were identified using the DESeq2 package in R 3.4.3 (CRAN). Genes with *p*adj<0.05 were considered differentially expressed. For principal component analysis (PCA), the 1000 most variable genes were used and data were mean-centered per gene. For pathway analysis with Toppgene Suite, genes with nominal *p*-value <0.05 were used.(26)

### GSEA

For GSEA, we screened published datasets in the GEO NCBI database repository containing expression profiling data on human ECs, for datasets related to angiogenesis, pregnancy, endothelial activation, stimulation with hormones or inflammatory mediators, shear stress, hypoxia, growth patterning and other factors considered relevant in preeclampsia. Gene sets of 50-500 genes were created from published gene sets based on differentially expressed genes with P<0.05. GSEA was performed by 1000 random permutations of the phenotypic subgroups to establish a null distribution of enrichment score against which a (normalized) enrichment score and nominal p-values were calculated.(27) Gene sets with p<0.05 were considered statistically significant.

### Multiplex immunoassay

Blood was collected in serum tubes within 4 hours before Caesarean section and spun down at 4000xg at 4°C. Serum was stored at −80°C until analysis. The multiplex immunoassay for 67 analytes was performed as described previously, measuring all analytes simultaneously in 50 μL of serum (xMAP; Luminex).(28) Heterophilic immunoglobulins were pre-absorbed from all samples with HeteroBlock (Omega Biologicals). Acquisition was performed with a Bio-Rad FlexMAP3D in combination with xPONENT software version 4.2 (Luminex). Data analysis was performed with Bioplex Manager 6.1.1 (Bio-Rad).

### Data analysis multiplex immunoassay

Multiplex data were analyzed using GraphPad Prism 7.0, SPSS Statistics 24 (IBM) and R 3.4.3 (CRAN). Out of ranges values on the lower end were imputed as 0.5x lowest measured value; out of ranges values on the upper end were imputed as 2x highest measured value. Analytes with more than 35% of measured values below the lower or above the upper limit of detection were excluded from the analyses (Granzyme B, Galectin-7, TRANCE-sRANKL, MIF, TNFa, IL-4, IL-1RA). For comparisons between two groups, the Mann-Whitney U test was used, with correction for multiple testing of all 60 analytes by Bonferroni or FDR as indicated where applicable. For principal component analysis (PCA) and heatmap analysis, data were mean-centered per analyte. Unsupervised hierarchical clustering was performed by Ward’s method with Euclidian distance. Random forest analysis was performed via http://www.metaboanalyst.ca/ with standard settings. Correlations were assessed by spearman rank correlation. Adjusted *p-*values <0.05 were considered significant.

## RESULTS

### Baseline characteristics

Maternal and pregnancy baseline characteristics for both the EC transcriptomics and the multiplex immunoassay are presented in Supplementary Table 2 and 3. Two patients with preeclampsia were excluded from the biomarker analyses due to cross-reactivity with the multiplex beads, which can lead to false-positive results. One additional preeclampsia case was excluded when histopathology did not confirm defective spiral artery remodeling. Baseline characteristics for both analyses were very similar. Women with preeclampsia were less often of white European descent and were more often obese, with both factors being known risk factors for the disorder. As expected, preeclampsia was associated with nulliparity, lower gestational age and birth weight.

### Placental bed endothelial cell transcriptomics

CD31^+^CD146^+^ ECs isolated from placental bed biopsied and sorted through flow cytometry-assisted cell sorting, were prepared for RNA-sequencing by Cel-Seq2 protocol. High expression of key endothelial genes such as Van Willebrand Factor, PECAM1 (CD31), MCAM (CD146) endoglin, claudin-5, CCL14, Tie1, CD34, and CTGF in addition to expression of 65 out of 72 endothelial-specific genes identified by Chi et al. confirmed endothelial cell identity (Supplementary Figure 1A).(29) The maternal origin of ECs was confirmed by high expression of the female-specific XIST gene and absent expression of the male-specific SRY gene in all samples. In addition to overlapping gene expression signatures between patients with preeclampsia and healthy pregnancies, we identified five differentially expressed genes (Figure 1A), including three significantly upregulated genes and two downregulated genes associated with preeclampsia (*p*adj<0.05; Figure 1B). Upregulated genes were prostaglandin D2 synthase (PTGDS), olfactomedin 1 (OFLM1) and IL-3 receptor subunit alpha (IL3RA). Downregulated genes were serine peptidase inhibitor Kazal type 5 (SPINK5) and sestrin 3 (SESN3). Three additional up- and three downregulated genes were identified with *p*adj<0.10 (Figure 1B), including nestin. Heatmap analysis comparing expression of the differentially expressed genes in preeclampsia with healthy pregnant controls is shown in Figure 1C. In order to perform pathway analysis of upregulated genes in preeclampsia we lowered the significance threshold (to a nominal *p*-value <0.05) to study potentially enriched pathways involved in preeclampsia pathogenesis. We identified 6 significantly enriched pathways within the transcriptional profile of the placental bed ECs (Figure 1D), related to both innate immune activation and platelet activation, suggesting inflammatory activation of ECs in preeclampsia. Comparison of EC transcriptional profiles at two sites within the uterus, i.e. the incision site and the placental bed site, between preeclampsia and healthy pregnancy, revealed partially overlapping gene signatures of upregulated and downregulated genes, including PTGDS and SPINK5 (Supplementary Figure 1B and 1C). This indicates that transcriptional changes occurring in preeclampsia may be present throughout the entire uterine vasculature, and not limited to the placental bed. Subsequent GSEA showed significant enrichment of genes upregulated in human umbilical vein endothelial cells (HUVECS) treated with plasma from patients with preeclampsia (figure 2A), indicating that circulating factors may partly induce the transcriptional phenotype observed in placental bed ECs from preeclamptic patients. In addition, GSEA showed significant enrichment of genes downregulated in HUVEC stimulated with VEGF or PlGF compared to their unstimulated counterparts, suggesting a VEGF- and PlGF deprived state in ECs of women with preeclampsia (figure 2B, 2C and 2D). Taken together, this analysis points towards significant enrichment of EC transcriptional changes which may be partly due to circulating factors involved in the pathogenesis of preeclampsia.

**Figure 1.**
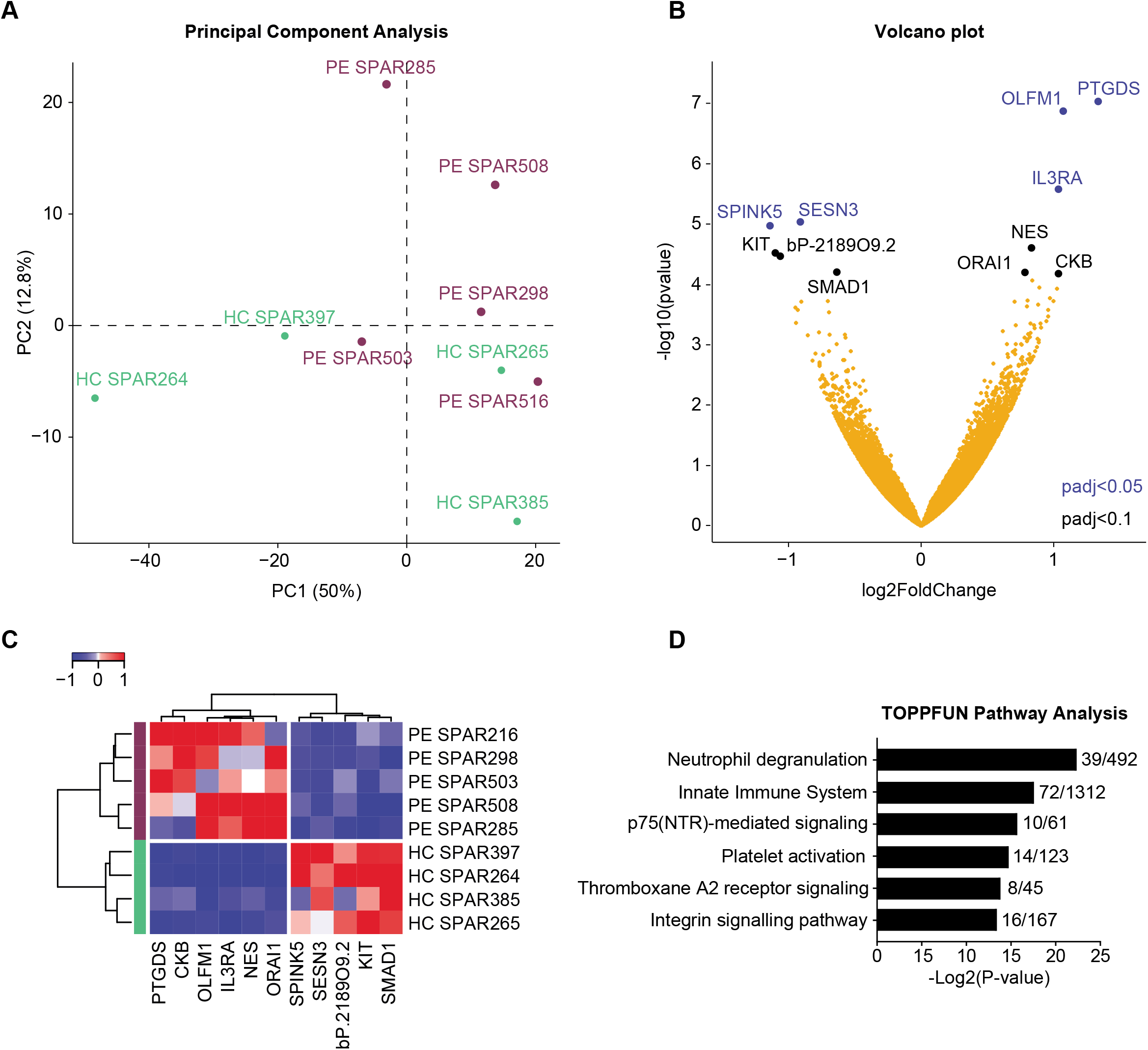
Transcriptomic profiling of spiral artery endothelial cells from the placental bed, comparing preeclampsia with FGR and healthy pregnancy. 2000 CD45^−^CD31^+^CD146^+^ endothelial cells were isolated from each biopsy by flow cytometry assisted cell sorting and RNA was sequenced by CEL-seq2 protocol. (**A**) Principal component analysis of preeclampsia cases and healthy controls using the 1000 genes with the highest variance (purple = preeclampsia, green = healthy controls). Genes were mean-centered. (**B**) Volcano plot showing differentially expressed genes with a *p*adj<0.05 (blue) and *p*adj<0.1 (black). (**C**) Heatmap of differentially expressed genes with a *p*adj<0.1. Genes were mean-centered and hierarchically clustered by Ward’s method and Euclidian distance. (**D**) Pathway analysis in ToppGene Suite on the 617 upregulated genes in preeclampsia compared to healthy pregnancy with a nominal *p*-value<0.05. Numbers indicate the number of overlapping upregulated genes in endothelial cells from preeclampsia samples, compared to the total known genes in the indicated pathway. *Abbreviations*: PE, preeclampsia (in this study early onset, in combination with fetal growth restriction); HC, healthy pregnancy; SPINK5, serine peptidase inhibitor Kazal type 5; SESN3,sestrin 3, KIT, KIT proto-oncogene receptor tyrosine kinase; SMAD1, Mothers against decapentaplegic homolog 1; PTGDS, prostaglandin D2 synthase; OLFM1, olfactomedin 1; IL3RA, interleukin 3 receptor subunit alpha; NES, nestin; ORAI1, Calcium release-activated calcium channel protein 1; CKB, creatine kinase B.

**Figure 2.**
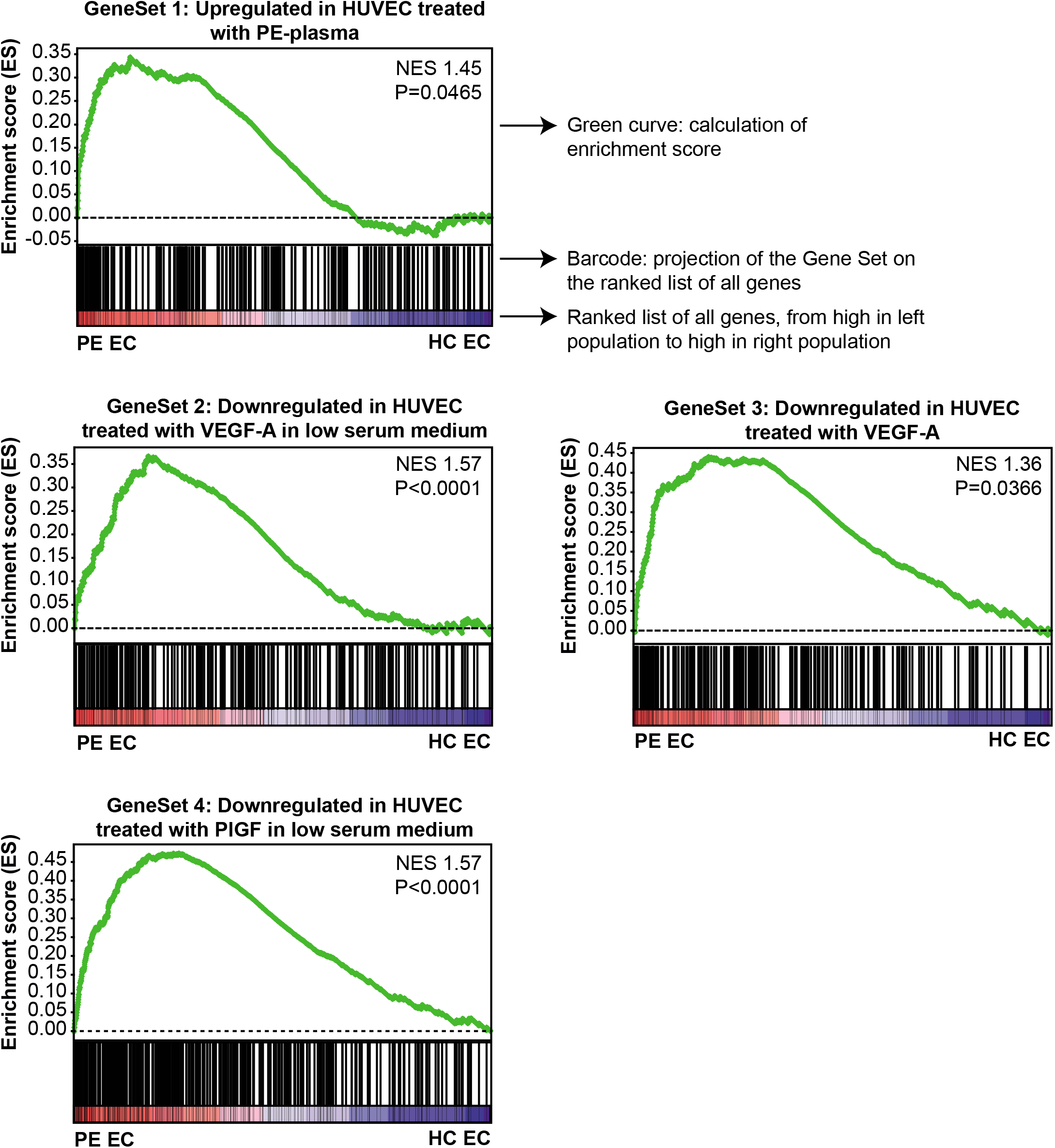
Gene set enrichment analysis (GSEA) showing significant enrichment of genes in published endothelial cell datasets. GSEA were run for all published datasets with endothelial cells (ECs) stimulated with factors relevant in preeclampsia. **(A)** GSEA with our preeclampsia cases showed significant enrichment for upregulated genes in HUVECS treated with PE plasma, and genes downregulated in HUVECs stimulated with VEGF **(B, C)** or PlGF **(D).**

### Markers of systemic inflammation and EC activation

Since the findings of GSEA analysis indicated that the presence or absence of circulating factors, known to be involved in preeclampsia, may have pathophysiological effects on the transcriptional profile of ECs, we performed biomarker profiling to further elucidate systemic disturbances of relevant proteins. To determine endothelial-related effects in preeclampsia we measured markers associated with inflammation, endothelial activation and endothelial dysfunction during the active disease state of preeclampsia patients compared with women with healthy pregnancy outcomes. Baseline characteristics are summarized in Supplementary Table 3. Using a comprehensive panel, we observed a clear and distinct biomarker signature associated with preeclampsia compared with normal pregnancy by principal component analysis (Figure 3A). A similar separation of groups was observed using unsupervised hierarchical clustering, with the exception of five individuals (Figure 3B). No clinical, histological or laboratory parameters could be identified to explain these exceptions, which merits further investigation. Analytes most contributing to separation of the two groups were identified as sFLT-1, endoglin and PlGF by random forest analysis, with an out-of-bag (OOB) error of 0.0 (Figure 3C). Accordingly, the sFLT-1/PlGF ratio was significantly increased in preeclampsia (Supplementary Figure 2), further confirming the preeclampsia phenotype. In addition to the sFlt-1/PlGF ratio, which is an established marker in preeclampsia, we also identified differences in leptin, and the acute phase reactant SAA-1 as highly discriminative factors, supporting the hypothesis that preeclampsia is associated with an pro-inflammatory phenotype (Figure 3D). This was further confirmed by increased levels of individual pro-inflammatory proteins IL-6, TNF-R1, and MIP-1β in preeclampsia, as compared to normal pregnant controls (Supplementary Table 4). Levels of endothelial activation markers sICAM-1 and E-selectin were also significantly higher in preeclampsia patients, further confirming the presence of systemic endothelial activation (Figure 3D, Supplementary Table 4).

**Figure 3.**
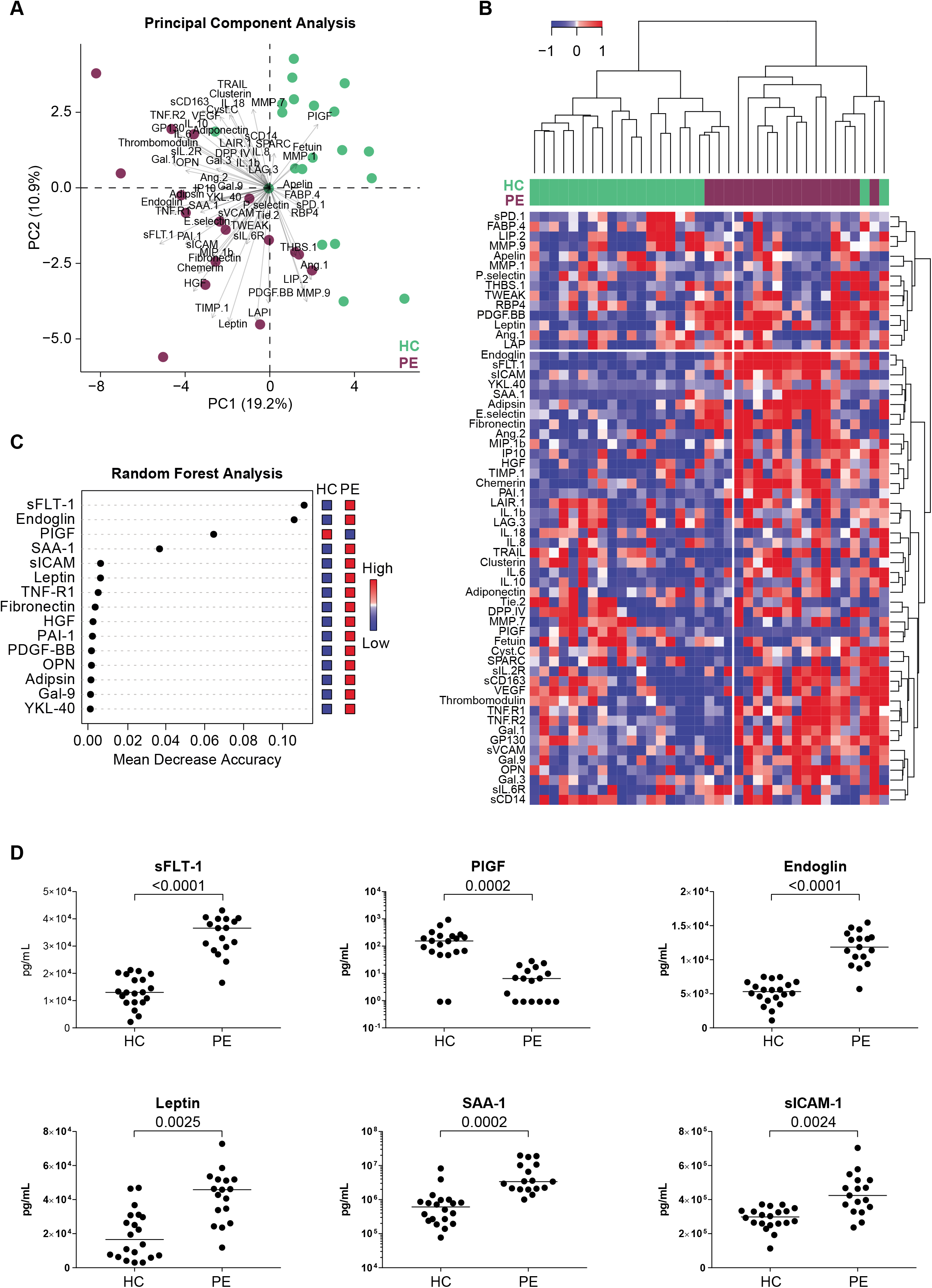
Systemic biomarker profiling of markers related to inflammation, endothelial activation and endothelial dysfunction, comparing preeclampsia with FGR and healthy pregnancy. Biomarkers were analyzed in serum by multiplex immunoassay. (**A**) Principal component analysis of preeclampsia cases and healthy controls using all 60 markers (purple = preeclampsia, green = healthy controls). Analytes were mean-centered. (**B**) Heatmap with hierarchical clustering of all 60 markers. Markers were mean-centered and patients were clustered by Ward’s method with Euclidian distance. (**C**) Random Forest analysis with 1000 trees yielding an out-of-bag error of 0.00 and showing the analytes most important for separation of preeclampsia and healthy groups. Analytes were mean-centered. (**D**) Scatter dot plots of sFLT-1, PlGF, Endoglin, Leptin, SAA-1, and sICAM-1. Line represents median; FDR with correction for multiple testing of 60 analytes is indicated. Mann-Whitney U test. *Abbreviations*: PE, preeclampsia (in this study early onset, in combination with fetal growth restriction); HC, healthy pregnancy; Multiplex Immunoassay abbreviations may be found in the Supplementary Tables.

To further explore heterogeneity within preeclampsia, unbiased analysis within the case group was performed. PCA and hierarchical clustering separated the case group into two clusters (Figure 4A and B). Strikingly, review of clinical and biochemical parameters revealed the presence of the clinically recognized severe preeclampsia phenotype of HELLP syndrome, in 6 out of 7 cases clustering separately. Analytes most contributing to this separation of preeclampsia with and without HELLP were identified as Ang-1, PDGF-BB, RPB4 and Apelin by random forest analysis with an OOB error of 0.294 (Figure 4C). All of these analytes were lower in patients with HELLP syndrome, which is likely due to the decreased platelet count in these women (*data not shown*). In addition, sFLT-1, SAA-1, adipsin, chemerin, and clusterin were higher in women with preeclampsia complicated by HELLP syndrome, although this effect was less clear after adjustment for multiple testing (Supplementary Table 5).

**Figure 4.**
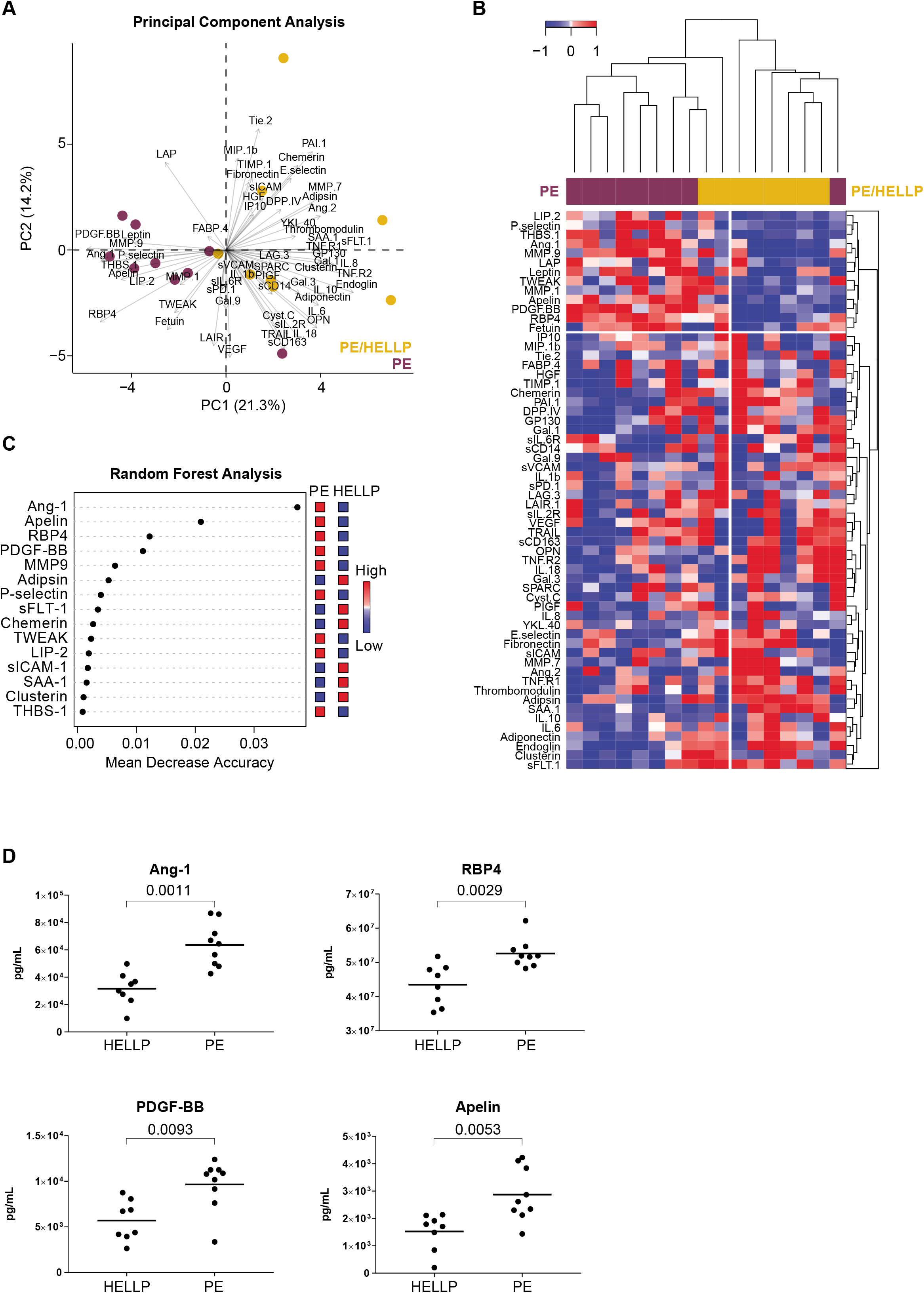
Systemic biomarker profiling of markers related to inflammation, endothelial activation and endothelial dysfunction within preeclampsia and a subgroup with HELLP syndrome. Biomarkers were analyzed in serum by multiplex immunoassay. (**A**) Principal component analysis of preeclampsia cases with and without HELLP using all 60 markers (purple = preeclampsia without HELLP, yellow = preeclampsia with HELLP). Analytes were mean-centered. (**B**) Heatmap with hierarchical clustering of all 60 markers. Markers were mean-centered and patients were clustered by Ward’s method with Euclidian distance. (**C**) Random Forest analysis with 1000 trees yielding an out-of-bag error of 0.294 and showing the analytes most important for separation of preeclampsia and healthy groups. Analytes were mean-centered. (**D**) Scatter dot plots of Ang-1, RBP4, PDGF-BB, and apelin. Line represents median; nominal p value without correction for multiple testing is indicated. Mann-Whitney U test. *Abbreviations*: PE, preeclampsia (in this study early onset, in combination with fetal growth restriction, without HELLP); HELLP, hemolysis, elevated liver enzymes, low platelets syndrome; Multiplex Immunoassay abbreviations may be found in the Supplementary Tables.

## DISCUSSION

In this study we used state of the art techniques to isolate ECs from the human placental bed to allow for transcriptomic profiling, comparing pregnancies complicated by preeclampsia with healthy pregnant controls. To our knowledge, our data provide the first transcriptomic analysis of human ECs isolated from this highly specialized vascular bed lying within the uterus at the maternal-fetal interface. We identified at least 5 differentially expressed genes (PTGDS, OLFM1, IL3RA, SPINK5 and SESN3) and pathways related to innate immune activation and platelet activation associated with preeclampsia. For two of these, PTGDS and SPINK5, differential expression was confirmed both at site of the uterus underlying the placenta, as well as at a second biopsy site elsewhere in the uterus, suggesting that some of the transcriptional changes observed in ECs from preeclampsia are confined to the placental bed while others are present in the entire uterus. While overall, transcriptomic profiling showed similarities between the two groups for individual genes, the pathway analysis identified distinct signatures associated with the disease. Consistent with other studies, EC activation and inflammation observed in the placental bed, could be found in the profiles of several key circulating markers of endothelial activation including the well-known markers sFLT1, endoglin and PlGF, as well as markers of inflammation including SAA-1 and leptin, further confirming the preeclampsia phenotype.

Our study provides proof-of-concept data that ECs derived from the placental bed in women with confirmed pathology of the spiral arteries underlying the placenta, indeed show signs of altered gene expression similar to endothelial activation and inflammation observed in other vascular beds. Although these findings need to be followed up in additional experiments, some of the differentially expressed genes have been associated with preeclampsia in previous studies, and may be considered as plausible candidates for involvement in the pathophysiology of the disease, as summarized below. PTGDS, upregulated in preeclampsia, catalyzes the conversion of prostaglandin H2 to prostaglandin D2 (PGD_2_) and is important for inhibition of platelet aggregation, relaxation and contraction of smooth muscle and reduction of vascular permeability.(30, 31) ECs are known to produce PTGDS under shear stress, which is likely present in the case of insufficient remodeling and high blood pressure in preeclampsia.(32–34) PGD_2_ may be involved in inflammation through recruitment of T helper (Th) type 2 cells.(35) Although many studies show increased Th1 type and decreased Th2 type immunity in preeclampsia, at least as many reports demonstrate the opposite.(13) PTGDS upregulation has been associated with uterine contraction and spontaneous preterm birth, which may be linked to preeclampsia due to belief of some that spontaneous preterm birth serves as an internal rescue mechanism aiming to protect both mother and baby from damaging effects of prolonged preeclamptic and growth restricted pregnancy.(36–41) Studies on olfactomedin 1 (Olfm-1) in the context of reproduction are limited. Human recombinant Olfm-1 suppresses the attachment of spheroids onto endometrial cells and downregulation of Olfm-1 during the receptive period may favor embryo attachment for successful implantation.(42) IL-3RA is a receptor for IL-3. IL-3 was shown to mediate positive signals for embryo implantation and to promote placental development and fetal growth.(43) IL-3RA receptor expression on EC increases migration of dendritic cells into tissues, which may regulate the Th1/Th2 balance within the decidua to maintain a Th2-dominant state, which is essential for maintenance of pregnancy.(44–46) A pathologic implication of high IL3RA expression in reproduction has not been reported. Nestin (NES), a type VI intermediate filament protein known to participate in remodeling of the cell, was borderline upregulated in preeclampsia. In animal models, NES upregulation characterizes vascular remodeling secondary to hypertension. In comparison to multiple other reproductive tissues, NES is most strongly expressed by ECs of newly vascularized tissues.(47, 48) In humans, urinary NES levels are significantly increased in preeclampsia patients and positively correlate with proteinuria, which is a key feature of the preeclampsia phenotype.(48) The SPINK5 gene, downregulated in ECs from preeclamptic patients, codes for the protein LEKT1, a serine protease inhibitor. Polymorphisms in SPINK5 have been associated with hypersensitivity of the immune system, especially in skin (atopy).(49) Other members of the SPINK family (i.e. SPINK1) have been shown to be highly up-regulated in decidua of recurrent pregnancy loss and to be predictive of preeclampsia, but no evidence is available for similar functions of SPINK5.(50) Finally, Sestrin 3, which was downregulated in preeclampsia in our study, reduces the levels of intracellular reactive oxygen species and is stress-induced.(51) It is required for normal regulation of blood glucose, insulin resistance, plays a role in lipid storage in obesity and is associated with increasing severity of coronary artery disease.(52, 53) SESN3 has not previously been investigated in reproduction.

Pathway analysis of the genes enriched in ECs from preeclampsia identified 6 upregulated pathways, involving the innate immune system and platelet activation. GSEA indicated that disturbances in circulating factors, especially related to angiogenesis, may contribute to the transcriptional changes observed in EC from preeclamptic patients. Systemic biomarker profiling by multiplex immunoassay confirmed immune activation and a disturbed balance between angiogenic and angiostatic factors in preeclampsia, which was most pronounced in patients with low platelets and elevated liver enzymes. These results indicate that inflammation and disturbed angiogenic signaling is not only present locally, in EC from the placental bed, but also systemically. Our findings are suggestive of the fact that this anti-angiogenic state plays a role in the generalized endothelial dysfunction key to preeclampsia, as well as contributes to the disturbed EC function at the placental bed, which may explain the increased susceptibility to endothelial cell erosion and lack of endothelial repair observed in defective spiral artery remodeling.(15)

Strengths of our study include the use of transcriptomics to investigate ECs from the human placental bed in patients with pregnancy complicated by severe preeclampsia and controls. This can only be performed on fresh tissue with 24/7 availability of dedicated clinicians and laboratory staff, as well as the use of state-of-the-art techniques for isolating very pure ECs from the placental bed with histologically confirmed defective spiral artery remodeling. Additionally, we confirmed the preeclampsia phenotype by extensive biomarker profiling to identify associated systemic markers of inflammation, endothelial dysfunction and soluble angiogenic factors. Limitations of this study include the limited availability of biopsy samples per group, which did not allow us to extensively confirm our findings in additional experiments e.g. immunohistochemistry or at the protein level. In addition, pregnancies of women with severe preeclampsia generally lead to delivery at an earlier gestational age than in women with planned Caesarean sections in the control group. In theory, the alterations in EC function attributed to preeclampsia may partly be influenced by this difference in gestational age. However, many of the markers we found to be discriminatory, confirmed the preeclampsia phenotype both at the EC, as well as at the systemic level.

In conclusion, we identified several candidate genes which were differentially expressed in endothelial cells from the placental bed that have a role in inflammatory response, vascular function and endothelial dysfunction, in addition to the known signs of generalized endothelial and inflammatory activation in preeclampsia. Our results underline the importance of maintaining vascular integrity by appropriate adaptation of ECs to the challenges of pregnancy, and contribute to the understanding of the impact of endothelial health on pregnancy outcome, as well as the similarities in the pathophysiology between preeclampsia and later-life arterial disease.

## Acknowledgements

We thank the multiplex core facility for their support.

---

**Supplementary Figure 1.**
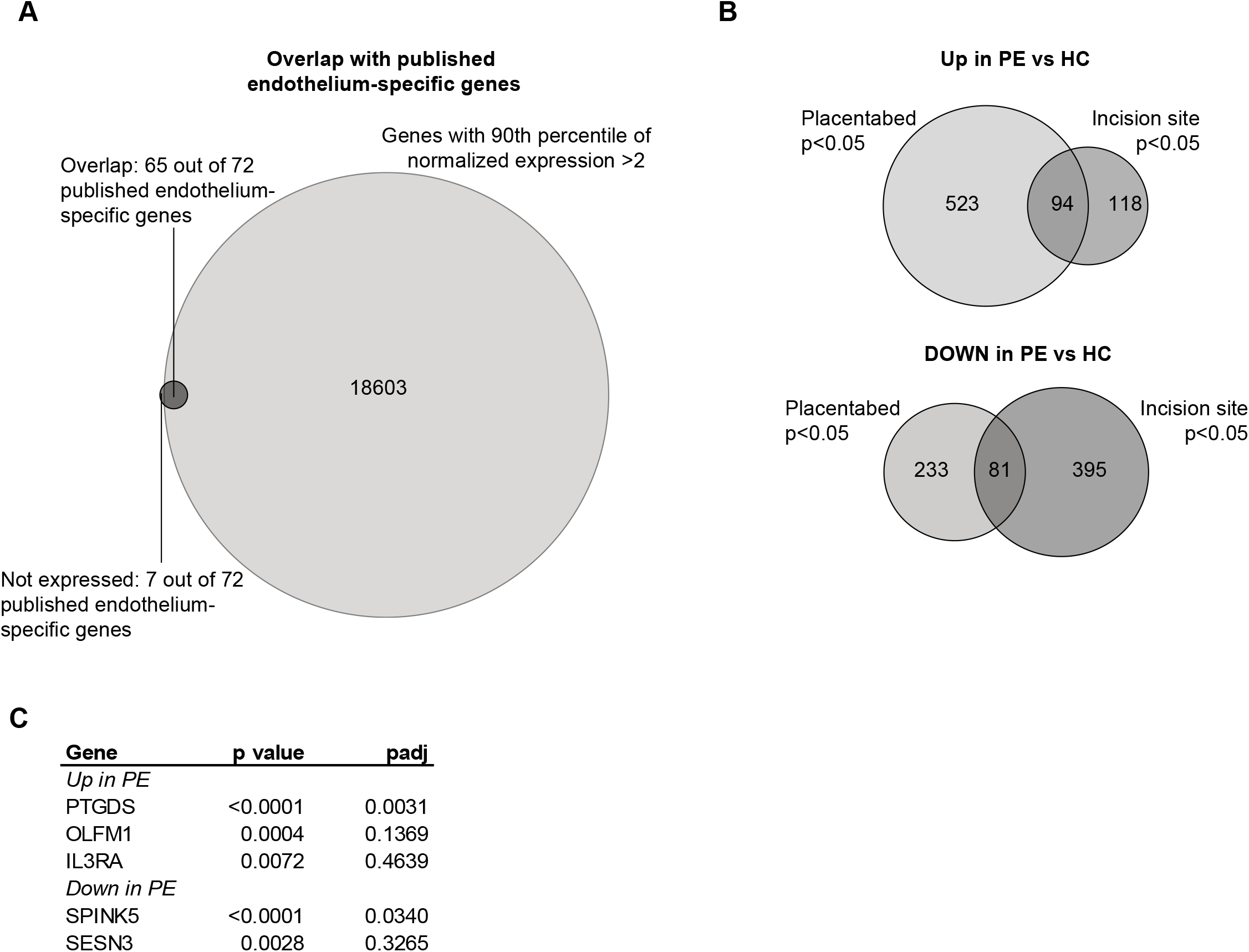
Endothelial identity and overlap of upregulated and downregulated genes in placental bed with incision site. **(A)** Overlap between 72 endothelium-specific genes published by Chi *et al*. and all genes with 90th percentile of normalized expression >2 in endothelial cells from the placental bed (irrespective of preeclampsia or healthy controls).^27^ **(B)** Overlap between upregulated and downregulated genes in preeclampsia compared to healthy pregnancies in placental bed and incision site, with a nominal *p*-value <0.05. (C) *P-*value and *p*adj of differential gene expression between preeclampsia and healthy controls at the incision site, for significantly differentially expressed genes with a *p*adj<0.05 in placental bed. *Abbreviations*: PE, preeclampsia (in this study early onset, in combination with fetal growth restriction); HC, healthy pregnancy; PTGDS, prostaglandin D2 synthase; OLFM1, olfactomedin 1; IL3RA, interleukin 3 receptor subunit alpha; SPINK5, serine peptidase inhibitor Kazal type 5; SESN3,sestrin 3.

**Supplementary Figure 2.**
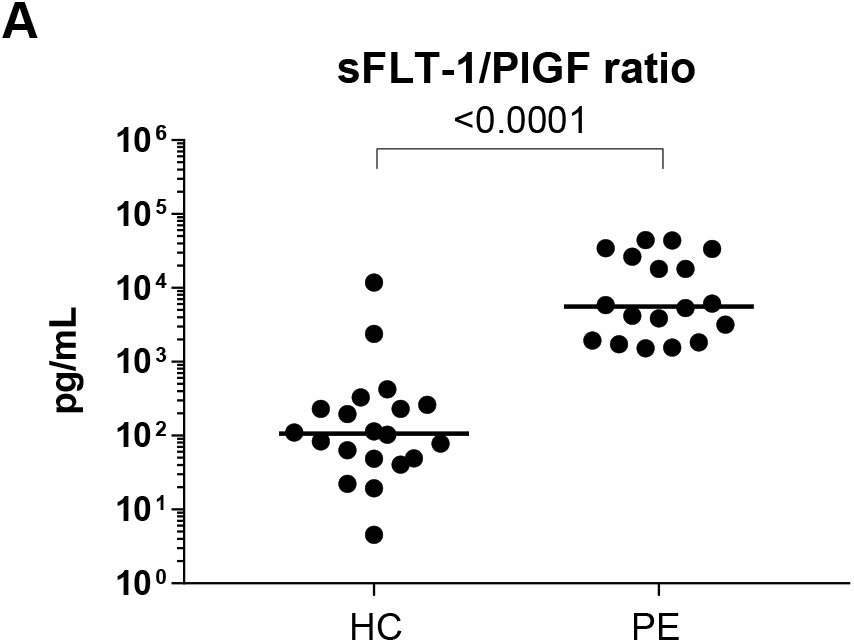
Ratio between serum sFLT-1 and PlGF in preeclampsia and healthy pregnancy. Scatter dot plots of sFLT-1/PlGF ratio. Line represents median; nominal *p-*value without correction for multiple testing is indicated and calculated by Mann-Whitney U test. *Abbreviations*: PE, preeclampsia (in this study early onset, in combination with fetal growth restriction); HC, healthy pregnancy; sFLT-1, soluble fms-like tyrosine kinase-1; sICAM, soluble Intercellular Adhesion Molecule; PlGF, Placental growth factor.

**Supplementary Table 1.**
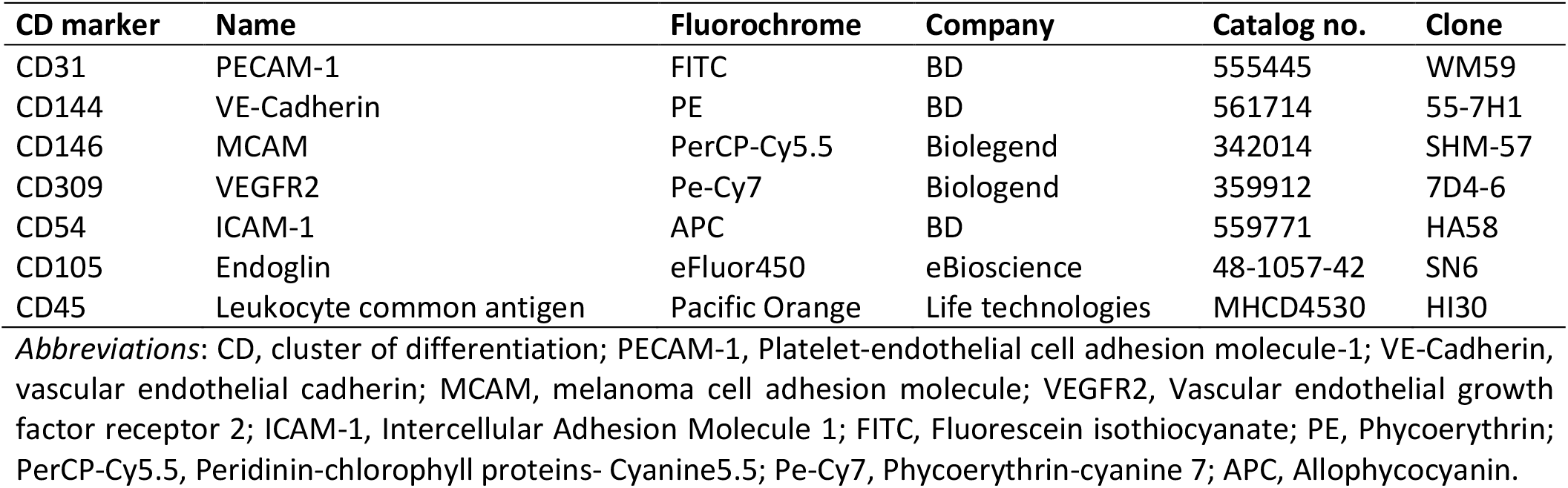
Antibodies used for flow cytometry assisted cell sorting.

**Supplementary Table 2.** Baseline characteristics for nulliparous women with preeclampsia and healthy pregnancy included for endothelial cell transcriptional profile comparison.

|  | Healthy control<br>*n*=4 | Preeclampsia<br>*n*=5 | *p*-value |
| --- | --- | --- | --- |
| **General characteristics** |  |  |  |
| Age (years) | 35.3 (5.3) | 28.2 (7.1) | 0.145 |
| White European (%) | 4 (100%) | 2 (40.0%) | 0.058 |
| BMI (kg/m<sup>2</sup>) | 26.6 (5.5) | 28.1 (5.6) | 0.699 |
| Obesity (%) | 2 (50.0%) | 2 (40.0%) | 0.764 |
| Smoking (%) | 0 (0.0%) | 0 (0.0%) | n/a |
| **General and obstetric history** |  |  |  |
| Pre-existent hypertension (%) | 0 (0.0%) | 0 (0.0%) | n/a |
| Pre-existent kidney disease (%) | 0 (0.0%) | 0 (0.0%) | n/a |
| Pre-existent Diabetes (%) | 0 (0.0%) | 0 (0.0%) | n/a |
| Nulliparity (%) | 4 (100.0%) | 5 (100.0%) | n/a |
| **Pregnancy characteristics** |  |  |  |
| Early onset (%) | n/a | 5 (100%) | n/a |
| FGR (%) | n/a | 5 (100%) | n/a |
| HELLP (%) | n/a | 2 (40%) | n/a |
| GDM with insulin use (%) | 0 (0.0%) | 0 (0.0%) | n/a |
| HPP (%) | 1 (25%) | 1 (20%) | 0.858 |
| **Maternal characteristics** |  |  |  |
| Systolic blood pressure (mmHg) | 122 (10) | 162 (22) | 0.010 |
| Diastolic blood pressure (mmHg) | 76 (8) | 101 (9) | 0.003 |
| Antihypertensive treatment oral (%) | 0 (0.0%) | 4 (80.0%) | 0.016 |
| Antihypertensive treatment iv (%) | 0 (0.0%) | 1 (20.0%) | 0.343 |
| Antepartum MgSO<sub>4</sub> iv (%) | 0 (0.0%) | 4 (80.0%) | 0.016 |
| Antepartum CCS (%) | 0 (0.0%) | 5 (100%) | 0.003 |
| **Neonatal characteristics** |  |  |  |
| GA at delivery (days) | 277 (7) | 216 (16) | <0.001 |
| Birthweight (grams) | 3529 (525) | 1135 (338) | <0.001 |
| Birthweight <3<sup>rd</sup> percentile (%) | 0 (0.0%) | 4 (80.0%) | <0.001 |
| Fetal sex, male (%) | 2 (50%) | 3 (60%) | 0.764 |

Values are presented as means (standard deviations) or as numbers (%). *P*-values were calculated by the $\chi^2$ test for continuous variables and Fisher's exact test for categorical variables. Abbreviations: BMI, Body Mass Index; Obesity was defined as an BMI >30kg/m<sup>2</sup>; FGR, fetal growth restriction; HELLP, Hemolysis Elevated Liver enzymes Low Platelets syndrome; GDM, gestational diabetes mellitus, HPP, hemorrhage post-partum, defined as blood loss >1000mL; iv, intravenous; MgSO<sub>4</sub>, magnesiumsulphate; CCS, corticosteroid treatment; GA, gestational age (days); n/a; not applicable.

**Supplementary Table 3.**
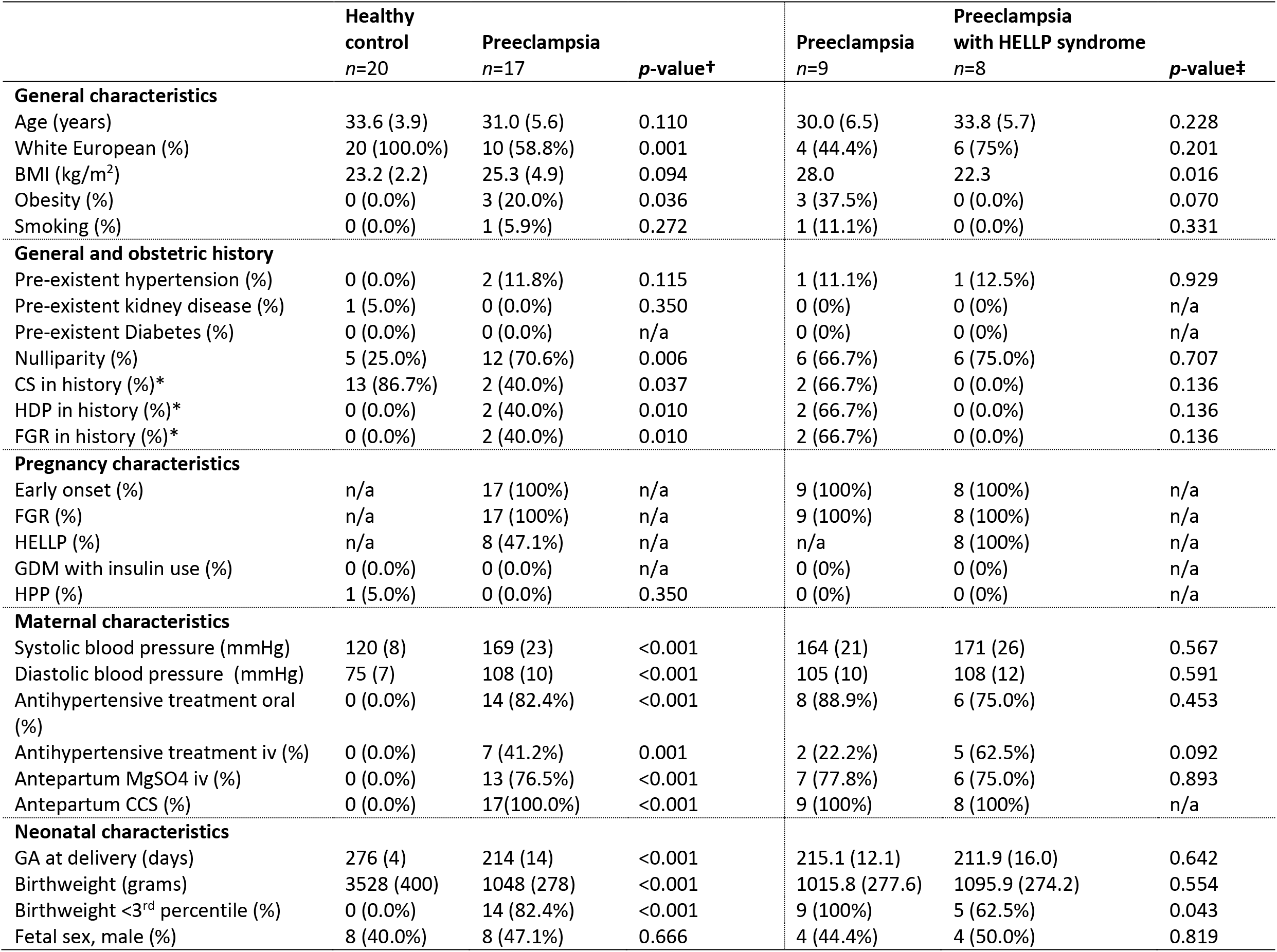
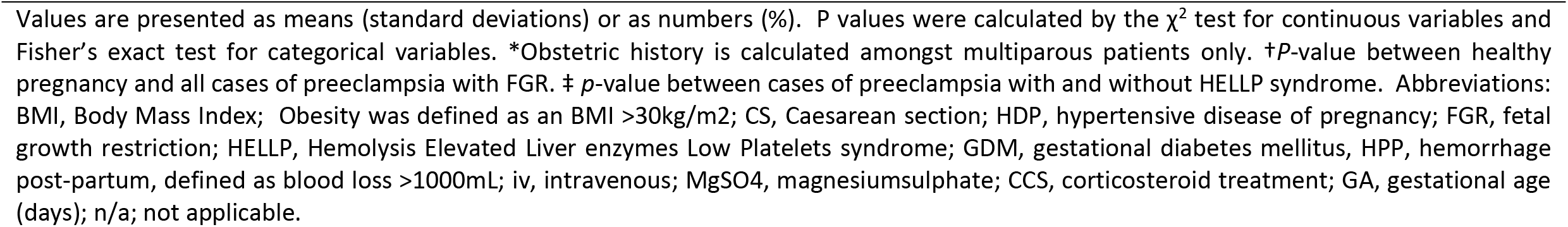
Maternal and pregnancy baseline characteristics in women with severe preeclampsia (with and without HELLP syndrome) and healthy pregnancy for the multiplex immunoassay comparison of systemic biomarkers for inflammation, endothelial activation and dysfunction.

|  | Healthy control<br>*n*=20 | Preeclampsia<br>*n*=17 | *p*-value† | Preeclampsia<br>*n*=9 | Preeclampsia with HELLP syndrome<br>*n*=8 | *p*-value‡ |
| --- | --- | --- | --- | --- | --- | --- |
| **General characteristics** |  |  |  |  |  |  |
| Age (years) | 33.6 (3.9) | 31.0 (5.6) | 0.110 | 30.0 (6.5) | 33.8 (5.7) | 0.228 |
| White European (%) | 20 (100.0%) | 10 (58.8%) | 0.001 | 4 (44.4%) | 6 (75%) | 0.201 |
| BMI (kg/m<sup>2</sup>) | 23.2 (2.2) | 25.3 (4.9) | 0.094 | 28.0 | 22.3 | 0.016 |
| Obesity (%) | 0 (0.0%) | 3 (20.0%) | 0.036 | 3 (37.5%) | 0 (0.0%) | 0.070 |
| Smoking (%) | 0 (0.0%) | 1 (5.9%) | 0.272 | 1 (11.1%) | 0 (0.0%) | 0.331 |
| **General and obstetric history** |  |  |  |  |  |  |
| Pre-existent hypertension (%) | 0 (0.0%) | 2 (11.8%) | 0.115 | 1 (11.1%) | 1 (12.5%) | 0.929 |
| Pre-existent kidney disease (%) | 1 (5.0%) | 0 (0.0%) | 0.350 | 0 (0%) | 0 (0%) | n/a |
| Pre-existent Diabetes (%) | 0 (0.0%) | 0 (0.0%) | n/a | 0 (0%) | 0 (0%) | n/a |
| Nulliparity (%) | 5 (25.0%) | 12 (70.6%) | 0.006 | 6 (66.7%) | 6 (75.0%) | 0.707 |
| CS in history (%)* | 13 (86.7%) | 2 (40.0%) | 0.037 | 2 (66.7%) | 0 (0.0%) | 0.136 |
| HDP in history (%)* | 0 (0.0%) | 2 (40.0%) | 0.010 | 2 (66.7%) | 0 (0.0%) | 0.136 |
| FGR in history (%)* | 0 (0.0%) | 2 (40.0%) | 0.010 | 2 (66.7%) | 0 (0.0%) | 0.136 |
| **Pregnancy characteristics** |  |  |  |  |  |  |
| Early onset (%) | n/a | 17 (100%) | n/a | 9 (100%) | 8 (100%) | n/a |
| FGR (%) | n/a | 17 (100%) | n/a | 9 (100%) | 8 (100%) | n/a |
| HELLP (%) | n/a | 8 (47.1%) | n/a | n/a | 8 (100%) | n/a |
| GDM with insulin use (%) | 0 (0.0%) | 0 (0.0%) | n/a | 0 (0%) | 0 (0%) | n/a |
| HPP (%) | 1 (5.0%) | 0 (0.0%) | 0.350 | 0 (0%) | 0 (0%) | n/a |
| **Maternal characteristics** |  |  |  |  |  |  |
| Systolic blood pressure (mmHg) | 120 (8) | 169 (23) | <0.001 | 164 (21) | 171 (26) | 0.567 |
| Diastolic blood pressure (mmHg) | 75 (7) | 108 (10) | <0.001 | 105 (10) | 108 (12) | 0.591 |
| Antihypertensive treatment oral (%) | 0 (0.0%) | 14 (82.4%) | <0.001 | 8 (88.9%) | 6 (75.0%) | 0.453 |
| Antihypertensive treatment iv (%) | 0 (0.0%) | 7 (41.2%) | 0.001 | 2 (22.2%) | 5 (62.5%) | 0.092 |
| Antepartum MgSO<sub>4</sub> iv (%) | 0 (0.0%) | 13 (76.5%) | <0.001 | 7 (77.8%) | 6 (75.0%) | 0.893 |
| Antepartum CCS (%) | 0 (0.0%) | 17(100.0%) | <0.001 | 9 (100%) | 8 (100%) | n/a |
| **Neonatal characteristics** |  |  |  |  |  |  |
| GA at delivery (days) | 276 (4) | 214 (14) | <0.001 | 215.1 (12.1) | 211.9 (16.0) | 0.642 |
| Birthweight (grams) | 3528 (400) | 1048 (278) | <0.001 | 1015.8 (277.6) | 1095.9 (274.2) | 0.554 |
| Birthweight <3<sup>rd</sup> percentile (%) | 0 (0.0%) | 14 (82.4%) | <0.001 | 9 (100%) | 5 (62.5%) | 0.043 |
| Fetal sex, male (%) | 8 (40.0%) | 8 (47.1%) | 0.666 | 4 (44.4%) | 4 (50.0%) | 0.819 |

Values are presented as means (standard deviations) or as numbers (%). P values were calculated by the $\chi^2$ test for continuous variables and Fisher's exact test for categorical variables. \*Obstetric history is calculated amongst multiparous patients only. †P-value between healthy pregnancy and all cases of preeclampsia with FGR. ‡ p-value between cases of preeclampsia with and without HELLP syndrome. Abbreviations: BMI, Body Mass Index; Obesity was defined as an BMI >30kg/m<sup>2</sup>; CS, Caesarean section; HDP, hypertensive disease of pregnancy; FGR, fetal growth restriction; HELLP, Hemolysis Elevated Liver enzymes Low Platelets syndrome; GDM, gestational diabetes mellitus, HPP, hemorrhage post-partum, defined as blood loss >1000mL; iv, intravenous; MgSO<sub>4</sub>, magnesiumsulphate; CCS, corticosteroid treatment; GA, gestational age (days); n/a; not applicable.

**Supplementary Table 4.**
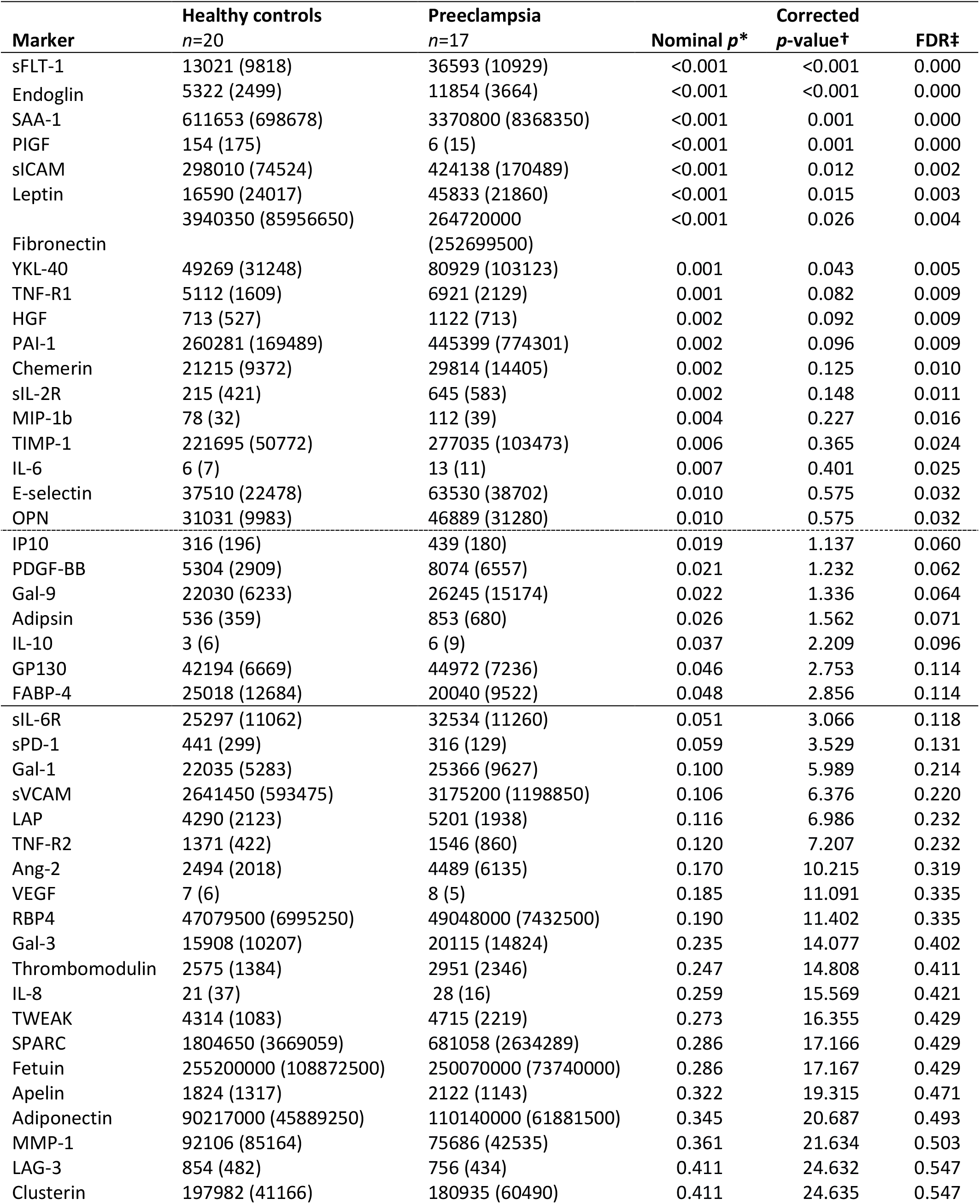
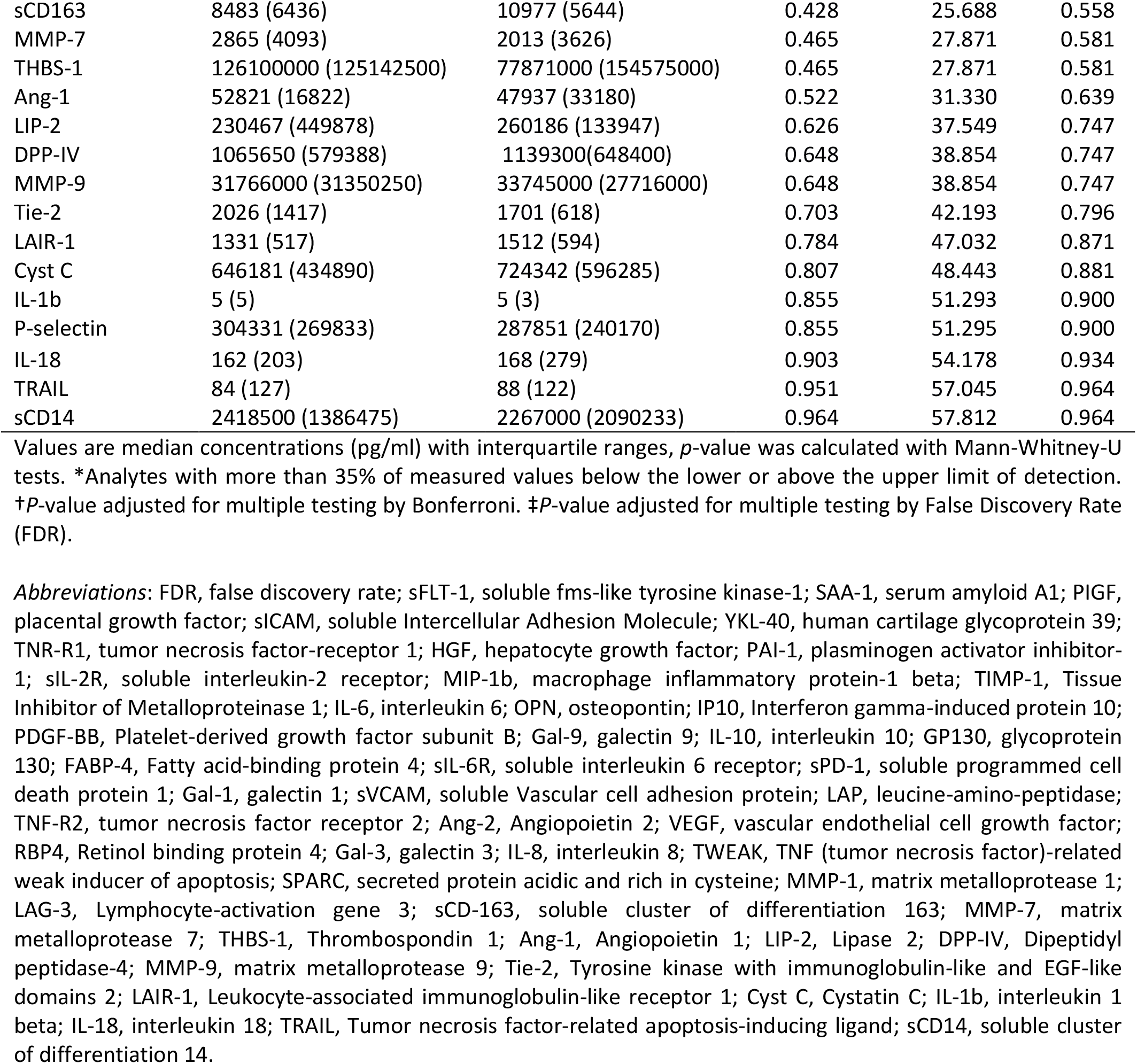
Individual marker comparison by multiplex immunoassay between preeclampsia and healthy pregnancy.

**Supplementary Table 5.** Individual marker comparison by multiplex immunoassay between women with preeclampsia with or without HELLP syndrome.

| Marker | Preeclampsia<br>*n*=9 | Preeclampsia<br>with HELLP syndrome<br>*n*=8 | Nominal *p* * | Corrected<br>*p*-value† | FDR‡ |
| --- | --- | --- | --- | --- | --- |
| Ang-1 | 64475 (30089) | 32536 (15733) | 0.001 | 0.064 | 0.064 |
| RBP4 | 51902000 (4672000) | 44490500 (11239000) | 0.003 | 0.171 | 0.086 |
| Apelin | 2613 (1764) | 1747 (1054) | 0.005 | 0.316 | 0.105 |
| PDGF-BB | 10789 (2871) | 5552 (3771) | 0.009 | 0.560 | 0.124 |
| sFLT-1 | 29677 (11717) | 40018 (6668) | 0.012 | 0.741 | 0.124 |
| P-selectin | 361210 (678401) | 199624 (212380) | 0.012 | 0.741 | 0.124 |
| LIP-2 | 308570 (235993) | 216820 (68172) | 0.016 | 0.969 | 0.138 |
| SAA-1 | 2152800 (1799600) | 10419500 (16176975) | 0.021 | 1.255 | 0.144 |
| Adipsin | 609 (652) | 992 (148) | 0.024 | 1.419 | 0.144 |
| Clusterin | 173716 (31083) | 205257 (83254) | 0.027 | 1.613 | 0.144 |
| MMP-9 | 38172000 (18494500) | 15480500 (27122425) | 0.027 | 1.613 | 0.144 |
| PAI-1 | 329237 (602442) | 702866 (873613) | 0.034 | 2.056 | 0.144 |
| IL-10 | 5 (7) | 10 (19) | 0.043 | 2.598 | 0.144 |
| LAG-3 | 693 (284) | 955 (343) | 0.043 | 2.598 | 0.144 |
| IL-8 | 26 (9) | 36 (22) | 0.043 | 2.598 | 0.144 |
| Endoglin | 10456 (3512) | 13188 (2584) | 0.043 | 2.598 | 0.144 |
| Chemerin | 24906 (14321) | 35253 (8072) | 0.043 | 2.598 | 0.144 |
| THBS-1 | 181170000<br>(222636500) | 61244500 (92863250) | 0.043 | 2.598 | 0.144 |
| E-selectin | 41700(35183) | 74198 (42960) | 0.054 | 3.258 | 0.171 |
| GP130 | 43643(14851) | 48886 (7553) | 0.068 | 4.050 | 0.193 |
| sICAM | 357172 (195616) | 466600 (181756) | 0.068 | 4.050 | 0.193 |
| Gal-1 | 22128 (8862) | 29001 (10229) | 0.083 | 4.996 | 0.217 |
| Ang-2 | 2365 (4026) | 5696 (7261) | 0.083 | 4.996 | 0.217 |
| TNF-R1 | 5749 (1907) | 7199 (2544) | 0.102 | 6.113 | 0.235 |
| TNF-R2 | 1412 (650) | 1910 (923) | 0.102 | 6.113 | 0.235 |
| LAP | 5420 (2165) | 4558 (1746) | 0.102 | 6.113 | 0.235 |
| Leptin | 51168 (18453) | 39824 (22068) | 0.112 | 6.728 | 0.249 |
| Fibronectin | 210150000<br>(218089825) | 329035000<br>(329784250) | 0.123 | 7.364 | 0.263 |
| TWEAK | 5407 (2120) | 4317 (735) | 0.149 | 8.935 | 0.298 |
| Tie-2 | 1699 (697) | 2150 (1201) | 0.149 | 8.935 | 0.298 |
| Thrombomodulin | 2526 (2500) | 3394 (2003) | 0.178 | 10.676 | 0.334 |
| DPP-IV | 1073200 (626920) | 1258550 (512500) | 0.178 | 10.676 | 0.334 |
| MMP-1 | 76927 (45814) | 67427 (64107) | 0.211 | 12.658 | 0.384 |
| Fetuin | 251270000<br>(57415000) | 213730000<br>(79657500) | 0.290 | 17.390 | 0.497 |
| TIMP-1 | 251428 (88968) | 281139 (75567) | 0.290 | 17.390 | 0.497 |
| FABP-4 | 22066 (11863) | 19132 (6272) | 0.336 | 20.155 | 0.530 |
| sVCAM | 3028100 (939800) | 3379500 (1542825) | 0.336 | 20.155 | 0.530 |
| Adiponectin | 91688000 (57801500) | 121705000<br>(66442500) | 0.336 | 20.155 | 0.530 |
| OPN | 44073 (18272) | 52769 (47517) | 0.386 | 23.189 | 0.580 |
| MMP-7 | 1999 (2219) | 2894 (5488) | 0.386 | 23.189 | 0.580 |
| SPARC | 1433700 (2914207) | 622578 (1582056) | 0.441 | 26.485 | 0.646 |
| IL-6 | 13 (10) | 15 (11) | 0.501 | 30.035 | 0.698 |
| HGF | 944 (854) | 1189 (612) | 0.501 | 30.035 | 0.698 |
| Gal-3 | 20115 (13396) | 23106 (23100) | 0.630 | 37.826 | 0.824 |
| sCD14 | 2543500 (2219787) | 2167300 (2068225) | 0.630 | 37.826 | 0.824 |
| IL-1b | 5 (4) | 5 (5) | 0.665 | 39.889 | 0.824 |
| VEGF | 8 (4) | 6 (6) | 0.700 | 41.998 | 0.824 |
| IL-18 | 168 (250) | 185 (327) | 0.700 | 42.019 | 0.824 |
| sIL-2R | 645 (642) | 717 (574) | 0.700 | 42.019 | 0.824 |
| Cyst C | 724342 (916234) | 727035 (514915) | 0.700 | 42.019 | 0.824 |
| Gal-9 | 23763 (16392) | 27282 (14181) | 0.700 | 42.019 | 0.824 |
| MIP-1b | 105 (39) | 114 (72) | 0.773 | 46.370 | 0.875 |
| sPD-1 | 326 (176) | 302 (54) | 0.773 | 46.370 | 0.875 |
| IP10 | 418 (164) | 440 (410) | 0.847 | 50.843 | 0.924 |
| sCD163 | 8599 (4445) | 11659 (8178) | 0.847 | 50.843 | 0.924 |
| PIGF | 6(14) | 4 (16) | 0.883 | 52.963 | 0.946 |
| sIL-6R | 32534(13135) | 32003 (10516) | 0.923 | 55.398 | 0.955 |
| LAIR-1 | 1512 (557) | 1482 (693) | 0.923 | 55.401 | 0.955 |
| TRAIL | 88 (101) | 65 (154) | 0.961 | 57.683 | 0.978 |
| YKL-40 | 80929 (74124) | 105167 (126816) | 1.000 | 60.000 | 1.000 |

Values are median concentrations (pg/ml) with interquartile ranges, *P*-value was calculated with Mann-Whitney-U tests. †*P*-value adjusted for multiple testing by Bonferroni. ‡*P*-value adjusted for multiple testing by False Discovery Rate (FDR).

## References

1. Powe CE, Levine RJ, Karumanchi SA. Preeclampsia, a disease of the maternal endothelium: the role of antiangiogenic factors and implications for later cardiovascular disease. Circulation. 2011;123(24):2856–69.

2. Duley L. The global impact of pre-eclampsia and eclampsia. Seminars in perinatology. 2009;33(3):130–7.

3. Kuklina EV, Ayala C, Callaghan WM. Hypertensive disorders and severe obstetric morbidity in the United States. Obstet Gynecol. 2009;113(6):1299–306.

4. Bellamy L, Casas JP, Hingorani AD, Williams DJ. Pre-eclampsia and risk of cardiovascular disease and cancer in later life: systematic review and meta-analysis. BMJ. 2007;335(7627):974.

5. de Jager SCA, Meeuwsen JAL, van Pijpen FM, Zoet GA, Barendrecht AD, Franx A, et al. Preeclampsia and coronary plaque erosion: Manifestations of endothelial dysfunction resulting in cardiovascular events in women. Eur J Pharmacol. 2017;816:129–37.

6. Brosens I, Pijnenborg R, Vercruysse L, Romero R. The “Great Obstetrical Syndromes” are associated with disorders of deep placentation. Am J Obstet Gynecol. 2011;204(3):193–201.

7. Fisher SJ. Why is placentation abnormal in preeclampsia? Am J Obstet Gynecol. 2015;213(4 Suppl):S115–22.

8. Lyall F, Robson SC, Bulmer JN. Spiral artery remodeling and trophoblast invasion in preeclampsia and fetal growth restriction: relationship to clinical outcome. Hypertension (Dallas, Tex : 1979). 2013;62(6):1046–54.

9. Staff AC, Dechend R, Pijnenborg R. Learning from the placenta: acute atherosis and vascular remodeling in preeclampsia-novel aspects for atherosclerosis and future cardiovascular health. Hypertension (Dallas, Tex : 1979). 2010;56(6):1026–34.

10. Staff AC, Johnsen GM, Dechend R, Redman CW. Preeclampsia and uteroplacental acute atherosis: immune and inflammatory factors. J Reprod Immunol. 2014;101-102:120–6.

11. Veerbeek JH, Brouwers L, Koster MP, Koenen SV, van Vliet EO, Nikkels PG, et al. Spiral artery remodeling and maternal cardiovascular risk: the spiral artery remodeling (SPAR) study. Journal of hypertension. 2016;34(8):1570–7.

12. Roberts JM. The perplexing pregnancy disorder preeclampsia: what next? Physiol Genomics. 2018;50(6):459–67.

13. Visser N, van Rijn BB, Rijkers GT, Franx A, Bruinse HW. Inflammatory changes in preeclampsia: current understanding of the maternal innate and adaptive immune response. Obstet Gynecol Surv. 2007;62(3):191–201.

14. Docheva N, Romero R, Chaemsaithong P, Tarca AL, Bhatti G, Pacora P, et al. The profiles of soluble adhesion molecules in the “great obstetrical syndromes”(). The journal of maternal-fetal & neonatal medicine : the official journal of the European Association of Perinatal Medicine, the Federation of Asia and Oceania Perinatal Societies, the International Society of Perinatal Obstet. 2018:1–24.

15. Ngene NC, Moodley J. Role of angiogenic factors in the pathogenesis and management of pre-eclampsia. International journal of gynaecology and obstetrics: the official organ of the International Federation of Gynaecology and Obstetrics. 2018;141(1):5–13.

16. Lopes van Balen VA, van Gansewinkel TAG, de Haas S, van Kuijk SMJ, van Drongelen J, Ghossein-Doha C, et al. Physiological adaptation of endothelial function to pregnancy: systematic review and meta-analysis. Ultrasound Obstet Gynecol. 2017;50(6):697–708.

17. McLaughlin K, Audette MC, Parker JD, Kingdom JC. Mechanisms and Clinical Significance of Endothelial Dysfunction in High-Risk Pregnancies. Can J Cardiol. 2018;34(4):371–80.

18. Alma LJ, Bokslag A, Maas A, Franx A, Paulus WJ, de Groot CJM. Shared biomarkers between female diastolic heart failure and pre-eclampsia: a systematic review and meta-analysis. ESC heart failure. 2017;4(2):88–98.

19. Roberts JM, Taylor RN, Musci TJ, Rodgers GM, Hubel CA, McLaughlin MK. Preeclampsia: an endothelial cell disorder. Am J Obstet Gynecol. 1989;161(5):1200–4.

20. Wu P, Haththotuwa R, Kwok CS, Babu A, Kotronias RA, Rushton C, et al. Preeclampsia and Future Cardiovascular Health: A Systematic Review and Meta-Analysis. Circulation Cardiovascular quality and outcomes. 2017;10(2).

21. Tranquilli AL, Dekker G, Magee L, Roberts J, Sibai BM, Steyn W, et al. The classification, diagnosis and management of the hypertensive disorders of pregnancy: A revised statement from the ISSHP. Pregnancy Hypertens. 2014;4(2):97–104.

22. Tranquilli AL, Brown MA, Zeeman GG, Dekker G, Sibai BM. The definition of severe and early-onset preeclampsia. Statements from the International Society for the Study of Hypertension in Pregnancy (ISSHP). Pregnancy Hypertens. 2013;3(1):44–7.

23. Terwisscha van Scheltinga JA, Scherjon SA, van Dillen J. NVOG-richtlijn Foetale groeirestrictie (FGR) Online: Nederlandse Vereniging voor Obstetrie en Gynaecologie; 2017 [Dutch guideline for fetal growth restriction (FGR)]. Available from: https://www.nvog.nl/wp-content/uploads/2017/12/Foetate-groeirestricie-FGR-15-09-2017.pdf.

24. Hashimshony T, Senderovich N, Avital G, Klochendler A, de Leeuw Y, Anavy L, et al. CEL-Seq2: sensitive highly-multiplexed single-cell RNA-Seq. Genome Biol. 2016;17:77.

25. Li H, Durbin R. Fast and accurate long-read alignment with Burrows-Wheeler transform. Bioinformatics. 2010;26(5):589–95.

26. Chen J, Bardes EE, Aronow BJ, Jegga AG. ToppGene Suite for gene list enrichment analysis and candidate gene prioritization. Nucleic Acids Res. 2009;37(Web Server issue):W305–11.

27. Subramanian A, Tamayo P, Mootha VK, Mukherjee S, Ebert BL, Gillette MA, et al. Gene set enrichment analysis: a knowledge-based approach for interpreting genome-wide expression profiles. Proc Natl Acad Sci U S A. 2005;102(43):15545–50.

28. Scholman RC, Giovannone B, Hiddingh S, Meerding JM, Malvar Fernandez B, van Dijk MEA, et al. Effect of anticoagulants on 162 circulating immune related proteins in healthy subjects. Cytokine. 2018;106:114–24.

29. Chi JT, Chang HY, Haraldsen G, Jahnsen FL, Troyanskaya OG, Chang DS, et al. Endothelial cell diversity revealed by global expression profiling. Proc Natl Acad Sci U S A. 2003;100(19):10623–8.

30. Lichtenstein SH, Carvell GE, Simons DJ. Responses of rat trigeminal ganglion neurons to movements of vibrissae in different directions. Somatosens Mot Res. 1990;7(1):47–65.

31. Omori K, Morikawa T, Kunita A, Nakamura T, Aritake K, Urade Y, et al. Lipocalin-type prostaglandin D synthase-derived PGD2 attenuates malignant properties of tumor endothelial cells. J Pathol. 2018;244(1):84–96.

32. Burton GJ, Woods AW, Jauniaux E, Kingdom JC. Rheological and physiological consequences of conversion of the maternal spiral arteries for uteroplacental blood flow during human pregnancy. Placenta. 2009;30(6):473–82.

33. Miyagi M, Miwa Y, Takahashi-Yanaga F, Morimoto S, Sasaguri T. Activator protein-1 mediates shear stress-induced prostaglandin d synthase gene expression in vascular endothelial cells. Arterioscler Thromb Vasc Biol. 2005;25(5):970–5.

34. Taba Y, Sasaguri T, Miyagi M, Abumiya T, Miwa Y, Ikeda T, et al. Fluid shear stress induces lipocalin-type prostaglandin D(2) synthase expression in vascular endothelial cells. Circ Res. 2000;86(9):967–73.

35. Saito S, Tsuda H, Michimata T. Prostaglandin D2 and reproduction. Am J Reprod Immunol. 2002;47(5):295–302.

36. Connealy BD, Carreno CA, Kase BA, Hart LA, Blackwell SC, Sibai BM. A history of prior preeclampsia as a risk factor for preterm birth. Am J Perinatol. 2014;31(6):483–8.

37. Enquobahrie DA, Williams MA, Qiu C, Muhie SY, Slentz-Kesler K, Ge Z, et al. Early pregnancy peripheral blood gene expression and risk of preterm delivery: a nested case control study. BMC Pregnancy Childbirth. 2009;9:56.

38. Kumar S, Palaia T, Hall CE, Ragolia L. Role of Lipocalin-type prostaglandin D2 synthase (L-PGDS) and its metabolite, prostaglandin D2, in preterm birth. Prostaglandins Other Lipid Mediat. 2015;118-119:28–33.

39. Liu B, Yang J, Luo W, Zhang Y, Li J, Li H, et al. Prostaglandin D2 is the major cyclooxygenase-1-derived product in prepartum mouse uteri where it mediates an enhanced in vitro myometrial contraction. Eur J Pharmacol. 2017;813:140–6.

40. Shiki Y, Shimoya K, Tokugawa Y, Kimura T, Koyama M, Azuma C, et al. Changes of lipocalin-type prostaglandin D synthase level during pregnancy. J Obstet Gynaecol Res. 2004;30(1):65–70.

41. Wikstrom AK, Stephansson O, Cnattingius S. Previous preeclampsia and risks of adverse outcomes in subsequent nonpreeclamptic pregnancies. Am J Obstet Gynecol. 2011;204(2):148 e1–6.

42. Kodithuwakku SP, Ng PY, Liu Y, Ng EH, Yeung WS, Ho PC, et al. Hormonal regulation of endometrial olfactomedin expression and its suppressive effect on spheroid attachment onto endometrial epithelial cells. Hum Reprod. 2011;26(1):167–75.

43. Fishman P, Falach-Vaknine E, Zigelman R, Bakimer R, Sredni B, Djaldetti M, et al. Prevention of fetal loss in experimental antiphospholipid syndrome by in vivo administration of recombinant interleukin-3. J Clin Invest. 1993;91(4):1834–7.

44. de la Rosa G, Longo N, Rodriguez-Fernandez JL, Puig-Kroger A, Pineda A, Corbi AL, et al. Migration of human blood dendritic cells across endothelial cell monolayers: adhesion molecules and chemokines involved in subset-specific transmigration. J Leukoc Biol. 2003;73(5):639–49.

45. Lim LH, Burdick MM, Hudson SA, Mustafa FB, Konstantopoulos K, Bochner BS. Stimulation of human endothelium with IL-3 induces selective basophil accumulation in vitro. J Immunol. 2006;176(9):5346–53.

46. Miyazaki S, Tsuda H, Sakai M, Hori S, Sasaki Y, Futatani T, et al. Predominance of Th2-promoting dendritic cells in early human pregnancy decidua. J Leukoc Biol. 2003;74(4):514–22.

47. Mokry J, Nemecek S. Angiogenesis of extra- and intraembryonic blood vessels is associated with expression of nestin in endothelial cells. Folia Biol (Praha). 1998;44(5):155–61.

48. Yang X, Ding Y, Yang M, Yu L, Hu Y, Deng Y. Nestin Improves Preeclampsia-Like Symptoms by Inhibiting Activity of Cyclin-Dependent Kinase 5. Kidney Blood Press Res. 2018;43(2):616–27.

49. Hubiche T, Ged C, Benard A, Leaute-Labreze C, McElreavey K, de Verneuil H, et al. Analysis of SPINK 5, KLK 7 and FLG genotypes in a French atopic dermatitis cohort. Acta Derm Venereol. 2007;87(6):499–505.

50. Krieg SA, Fan X, Hong Y, Sang QX, Giaccia A, Westphal LM, et al. Global alteration in gene expression profiles of deciduas from women with idiopathic recurrent pregnancy loss. Mol Hum Reprod. 2012;18(9):442–50.

51. Rhee SG, Bae SH. The antioxidant function of sestrins is mediated by promotion of autophagic degradation of Keap1 and Nrf2 activation and by inhibition of mTORC1. Free Radic Biol Med. 2015;88(Pt B):205–11.

52. Nascimento EB, Osler ME, Zierath JR. Sestrin 3 regulation in type 2 diabetic patients and its influence on metabolism and differentiation in skeletal muscle. Am J Physiol Endocrinol Metab. 2013;305(11):E1408–14.

53. Ye J, Wang M, Xu Y, Liu J, Jiang H, Wang Z, et al. Sestrins increase in patients with coronary artery disease and associate with the severity of coronary stenosis. Clin Chim Acta. 2017;472:51–7.

